# Streamlined and quantitative detection of chimerism in mouse tissue using digital PCR

**DOI:** 10.1101/2020.11.04.368944

**Authors:** Fabian P. Suchy, Toshiya Nishimura, Adam C. Wilkinson, Maimi Higuchi, Joydeep Bhadury, Hiromitsu Nakauchi

**Author notes:** Correspondence (F.P.S.), (H.N.).

## Abstract

Animal chimeras are widely used for biomedical discoveries, from developmental biology to cancer research. However, the accurate quantitation of mixed cell types in chimeric and mosaic tissues has challenges. Here, we have developed and characterized a droplet digital PCR single-nucleotide discrimination assay to detect chimerism among albino and non-albino mouse strains. In addition, we have validated that this assay is compatible with crude lysate from most organs, drastically streamlining sample preparation. This chimerism detection assay has many additional advantages over existing methods including its robust nature, minimal technical bias, and ability to report the total number of cells in a prepared sample. Importantly, the concepts developed and discussed here are readily adapted to other genomic loci to accurately measure mixed cell populations in any tissue.

## INTRODUCTION

Chimeric animal models have broad and important applications in the bioscientific and medical community. For example, stem and progenitor cells are injected into animals to characterize engraftment and function(Wood et al., 2012), patient-derived xenografts are frequently used to explore tumor growth *in vivo*(Hidalgo et al., 2014), and most transgenic animals are generated via chimeric founders. More recently, systemic animal chimeras have even been created to explore developmental biology and organogenesis, with exciting applications such as generating transplantable human organs(Goto et al., 2019; Suchy and Nakauchi, 2017; Yamaguchi et al., 2017). Determining accurate levels of chimerism in these research models is necessary to assess experimental phenotypes and judge treatment efficacy. However, measuring chimerism can be challenging, particularly if chimerism is very low.

Flow-cytometry (FCM) is extensively used for precise quantitation of mixed cell populations. Prior to analysis by FCM, tissues must be dissociated into a single-cell suspension. Fluorescently labelled antibodies or transgenically-expressed fluorophores can then be used to identify specific cell populations when run through a flow-cytometer. In the context of animal chimeras, if the cells are significantly different (e.g., human and mouse), antibodies with high specificity are available to distinguish the two populations(Köhler and Milstein, 1975). For detecting chimerism in allogenic samples, constitutively-expressed fluorescent transgenes are frequently used for identification(Kobayashi et al., 2010). However, these approaches have multiple limitations. First, fluorescent transgenes often silence, and silencing can be biased by cell type(Meyer, 1995). Second, few antibodies are available to distinguish populations with high genetic similarity. Third, single-cell dissociation may be incomplete or result in cell death for specific tissues, thus imparting a dissociation bias. Although antibodies such as anti-mouse CD45.1 and CD45.2 have been designed to study differences in syngenic and allogenic strains(Morse, 1992), phenotypic differences have been recently observed between the C57BL/6 CD45.1 and CD45.2 strains(Waterstrat et al., 2010). Therefore, other quantification methods are needed that are resistant to silencing, have less technical bias, and can distinguish subtle differences between two similar cell populations.

Digital PCR (dPCR) is an analytical tool that can be used as an alternative to flow-cytometry for absolute quantification of chimerism, with many benefits. Since all cells are lysed and DNA extracted in preparation for dPCR, tissue samples can be frozen, fixed, or even sectioned. Thus, samples can be prepared and analyzed later. In addition, cell-lysis protocols are generally more powerful and have less tissue-specific biases than enzymatic cell-dissociation buffers(Mitra et al., 2013). This eliminates dissociation bias and greatly streamlines sample preparation. Regarding specificity, reactions can be prepared with primers and hydrolysis-probes that distinguish a single nucleotide difference. This enables the flexibility to quantify ratios of nearly identical cell types.

The dPCR detection strategy is more robust and quantitative than other PCR-based approaches. After sample preparation, each dPCR reaction is partitioned into thousands of microreactions, such that they contain a either a single copy of target DNA or no copy. This is conceptually similar to flow-cytometry during which cell-suspensions are hydrodynamically focused into single column and then partitioned into individual droplets. After thermocycling, the dPCR microreactions are scored as positive or negative (digital output) from which the precise concentration of target DNA can be discerned. In contrast to common PCR and qPCR methods, the quantification strategy for dPCR does not require consistent or efficient amplification. dPCR is therefore more resistant to inhibitors and results in absolute quantification without references, as opposed to the relative quantification used in qPCR.

Here we describe a novel method for analyzing chimerism across albino and non-albino mouse strains using a single-nucleotide discrimination (SND) droplet digital PCR assay (ddPCR is type of dPCR that uses microdroplets to partition the PCR reaction)(Hindson et al., 2011; Whale et al., 2016). The assay is extensively characterized with numerous mouse strains, dilution series, and crude lysates from various tissues. Although the assays developed here have direct utility, concepts developed and discussed in this manuscript can be applied to any cell-type to robustly quantify cellular abundance based on a single-nucleotide difference.

## RESULTS

### Development and optimization

Many common laboratory mouse strains carry a homozygous guanine to cytosine substitution in the tyrosinase (*Tyr*) gene which causes a C103S amino acid change and results in a failure to produce melanin and the albino phenotype(Jackson and Bennett, 1990). We designed a SND-ddPCR assay to differentiate and quantitate the abundance of the wild-type (*Tyr*^*WT*^) and albino (*Tyr*^*alb*^) tyrosinase allele. The assay utilized forward/reverse primers that flank C103 to amplify a 100 bp region, a 6-carboxyfluorescein (FAM) labelled hydrolysis probe that matches the albino allele, and a hexachloro-fluorescein (HEX) labelled hydrolysis probe that matches wild-type allele (Figure 1a). As these probes differ only by a single nucleotide and are required to be multiplexed in each reaction, incorrect annealing to the mismatched amplicon is inevitable. As depicted in Figure 1a (right), optimizing the ddPCR annealing temperature results in minimal mismatch annealing, while a match results in maximal annealing.

**Figure 1 legend.**
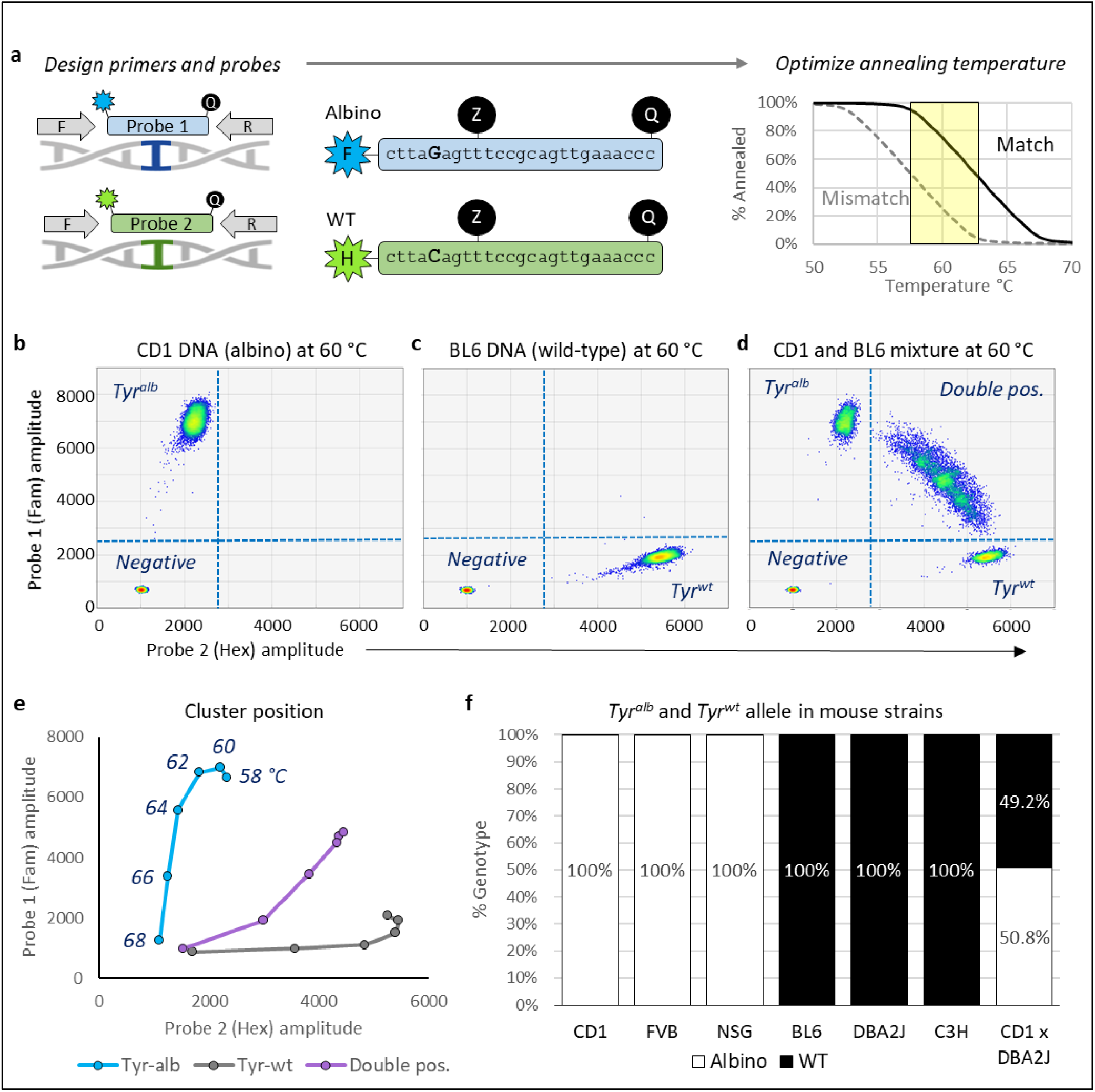
SND-ddPCR assay development and optimization. **a**. Schematic representing primer/probe annealing and optimization strategy for the SND-ddPCR assay. Left: F and R indicate shared forward and reverse primers respectively. Probes 1 and 2 are hydrolysis probes with FAM (blue) or HEX (green) fluorophores on the 5’ (left) end, and a quencher on the 3’ (right) end. Middle: Sequence of probes used to detect albino or WT allele. Internal ZEN quencher (Z) was used to reduce background. Bold letter indicates single-base difference between probes. Right: Theoretical melt curve for hydrolysis probes. Solid line represents probe annealing when all bases are matched to the target; dashed line represents annealing with a single base mismatch. The ideal temperature range for SND assays is highlighted in yellow, which results in maximum separation between matched and mismatched annealing. **b-d**. 2D display of SND-ddPCR results. Clusters composed of positive or negative microreactions are shown as heat map in each quadrant. CD1, BL6 and CD1+BL6 mixture are shown in b, c, and d respectively. **e**. 2D display showing centroid of the Tyr^Alb^, Tyr^WT^, and double positive cluster when analyzed at various annealing temperatures. The highest temperature is shown at the bottom left, and the lower temperatures move along the respective line upward and/or rightward. **f**. Frequency of Tyr^Alb^ and Tyr^WT^ allele detected in various mouse strains. CD1xDBA2J is F1 hybrid.

Column-purified CD-1 (CD1, albino) and C57BL/6 (BL6, wild-type) genomic DNA were first analyzed in different reactions with an annealing temperature of 60 °C (Figure 1b and c). The assay worked as expected: a negative cluster that contained droplets without *Tyr* was in the lower left quadrant; droplets containing *Tyr*^*Alb*^ from the CD1 mouse strain were detected with the FAM probe in the upper left quadrant (Figure 1b); and droplets containing *Tyr*^*WT*^ from the BL6 mouse strain were detected with the HEX probe in the lower right quadrant (Figure 1c). CD1 and BL6 DNA were then mixed at a 1:1 ratio and characterized by ddPCR (Figure 1d). This resulted in an arc-shaped cluster dispersed throughout the upper right quadrant, which contains droplets with at least one copy of both *Tyr*^*Alb*^ and *Tyr*^*WT*^. This cluster shape is typical of SND-ddPCR assays and is a result of *partition specific competition*(Whale et al., 2016). However, high concentrations of *Tyr* could lead to broadening of the double positive cluster, hindering the resolution from other clusters. Since counting the droplets in each of the four quadrants is critical to the accuracy of this assay, an upper limit of ~8 ng/ul mouse genomic DNA (~3000 copies of *Tyr*/ul) was selected for future reactions.

Figure 1b indicates that the HEX probe was mismatch-annealing (i.e., cross-reacting) with *Tyr*^*Alb*^, as indicated by the slight rightward shift of the *Tyr*^*Alb*^ cluster when compared to the negative cluster. The reciprocal is true of the FAM probe with *Tyr*^*WT*^ (Figure 1c). Therefore, the reactions were rerun at multiple annealing temperatures to find conditions optimal for cluster separation (Figure 1e). Although there was less mismatch-annealing ≥64 °C, the match-annealing started to drop. Maximum cluster separation was achieved at 60-64 °C. All further reactions were run at 60 °C.

### Characterization of common mouse strains

Having optimized the SND-ddPCR assay for detection of the *Tyr*^*Alb*^ and *Tyr*^*WT*^ allele, we used it to genotype different mouse strains. Genomic DNA was column purified from both white mice (CD1, FVB/NJ, NSG) and black/brown mouse mice (BL6, DBA/2J, C3H). The white mice only had the albino allele and the black/brown mice only had the WT allele (Figure 1f). To further validate our system, we generated *Tyr*^*Alb/WT*^ hybrid mice by crossing a CD1 male with a DBA/2J female. As expected, DNA analyzed from this hybrid strain showed a 1:1 ratio of *Tyr*^*Alb*^:*Tyr*^*WT*^. These data suggest that the SND-ddPCR assay could be used to detect chimerism between any of the white and brown/black mice analyzed.

### Dilution series: Accuracy and precision

To validate the accuracy and precision of the SND-ddPCR assay at various allele ratios and concentrations, a dilution matrix was prepared (Figure 2a). The chimerism varied from 20% to 0.03%, and the total concentration ranged from 2670 to 8.3 copies/ul (8 ng/ul to 25 pg/ul respectively). The assay could accurately report concentration and %chimerism at all concentrations above 0.5 copies/ul (Figure 2b; Supplementary Figure 1). Importantly, the concentration of the high abundance allele did not significantly influence the accuracy of detecting the low abundance allele. Precision also decreased significantly below 0.5 copies/ul as seen by a marked increase in variation (Figure 2c). Precise detection of the low abundance allele was not influenced by the quantity of the high abundance allele. The measured variation closely matched the theoretical variation in ddPCR at low concentrations due to random sampling distribution (R^2^ = 0.940, Figure 2c). Collectively, two important conclusions can be drawn: (1) accuracy and precision of the low abundance allele are not influenced by the high abundance allele, thus loading more DNA will increase the sensitivity; and (2) since the measured variation matches Poisson variation, confidence intervals can be calculated from Poisson distributions.

**Figure 2 legend.**
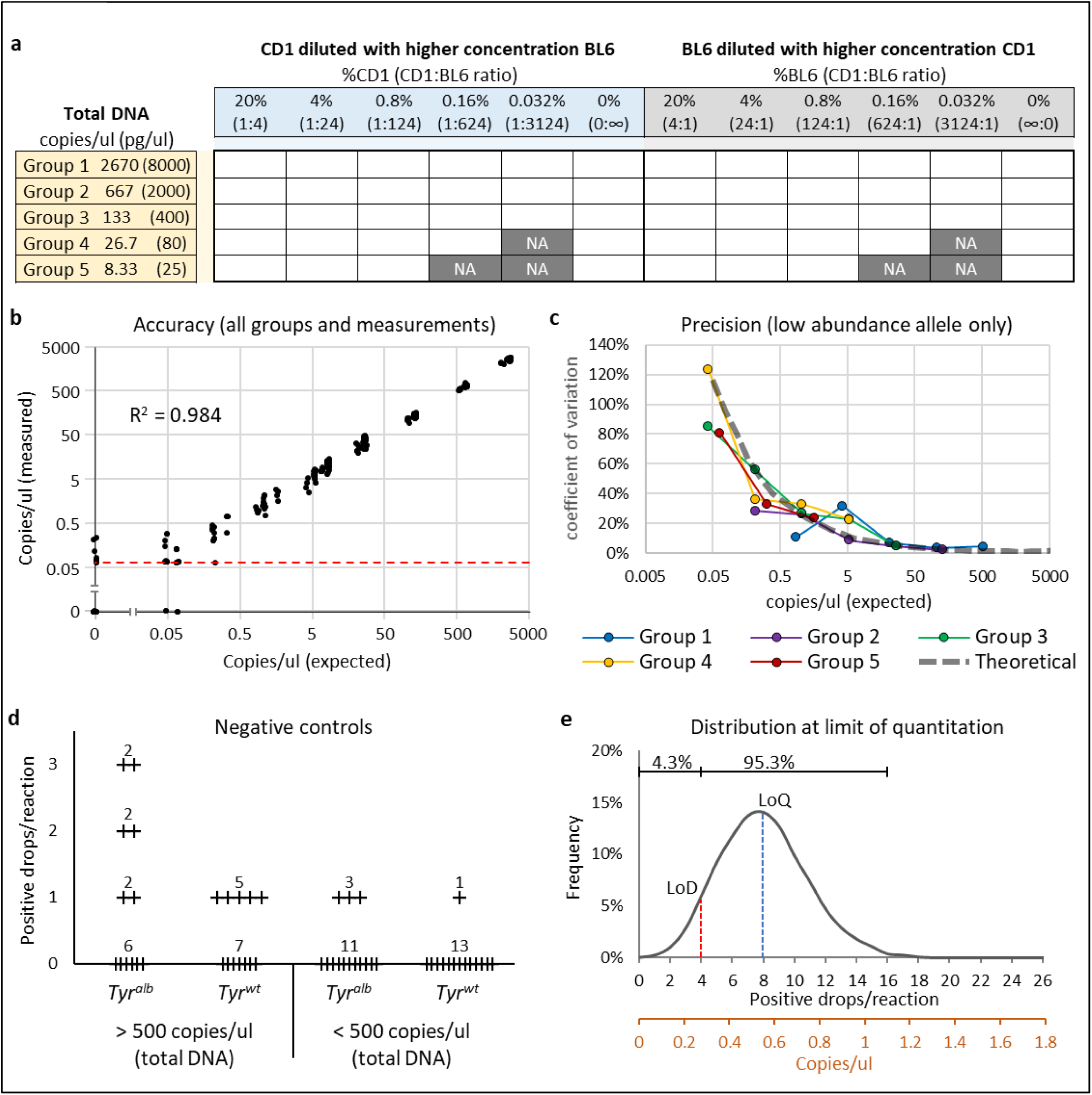
Accuracy and precision at various dilutions. **a**. Dilution table of CD1 and BL6 genomic DNA mixtures. Total DNA concentration was the same for each group. Dark gray boxes labeled NA were not analyzed because the concentration of low-abundance DNA was less than 0.5 molecules per reaction. **b**. Combined Group 1-5 concentration measurements compared to expected concentration; axes are log-scaled. 5% random error added to each point to aid in distinguishing overlapped datapoints. Red dashed line indicates the concentration at which there is only a single positive droplet per reaction (single molecule detection limit). R^2^ calculated from log-transformed data comparing measured and expected values (0.005 added to values with 0). **c**. Coeffecient of variation (CoV, standard deviation divided by mean) calculated for low-abundance (diluted) allele in Group 1-5 (n = 4 for each point: combined low-abundance CD1 and BL6 measurements, and two replicates of each). Gray dashed bar indicates theoretical CoV from Poisson distribution. R^2^ calculated from theoretical CoV compared to measured. **d**. Number of false positive droplets in negative controls. Analysis of 52 samples that contained just CD1 DNA, just BL6 DNA or no DNA. Combined data set from dilution series measurements and temperature optimization measurements (58-64 °C). Samples with just CD1 DNA used as negative controls for BL6; samples with just BL6 DNA used as negative controls for samples with CD1. **e**. Theoretical Poisson distribution of measured concentration when a sample is prepared at the LoQ (8 molecules per reaction). Black x-axis shows concentration as positive drops/reaction; brown x-axis shows concentration as copies/ul. >95% of the distribution lies with a factor of two from the LoQ (4-16). <5% of the distribution lies below the set LoD (<4 molecules per reaction). Each reaction assumed to have 17000 droplets.

### LoD and LoQ

Understanding the limits and confidence intervals of analytical tools is critical to interpreting results. Therefore, we next established a lower limit of detection (LoD) and lower limit of quantitation (LoQ). The false positive rate was considered to set an appropriate LoD. Although the theoretical LoD of PCR is a single molecule, we had up to three false positive droplets in multiple reactions (Figure 2d). Of the 52 negative controls, 15 (29%) detected at least one false positive droplet. The false positive rate slightly varied with overall concentration and was more prevalent for *Tyr*^*Alb*^. Regardless, a conservative LoD was set at 0.28 copies/ul (4 positive drops per reaction), which results in a stringent 0% false positive rate for this data set. The LoQ was set in a less stringent manner, such that there was ≥95% confidence that the measured value was at least within a factor of two from the true value. Poisson distribution shows that this is achieved at ≥0.55 copies/ul (8 positive drops per reaction, Figure 2e). At the LoQ, there is only a 4.3% chance that the assay would report a value below the LoD, resulting in a false negative.

### Blood chimerism: ddPCR vs flow cytometry

Up to this point, all analyses were performed with column-purified samples of genomic DNA that had been mixed to simulate chimeric samples. It is important to determine if the SND-ddPCR assay can accurately report chimerism when DNA is extracted from actual chimeric tissues. Blood is a convenient organ with which to validate this assay because it can be mixed at various ratios, readily analyzed by FCM, and directly compared to ddPCR without concern of heterogeneity or cell-dissociation bias. Since CD1 and BL6 mice express CD45.1 and CD45.2 respectively, fluorescently labelled antibodies can distinguish the two populations (Figure 3a). Blood from CD1 and BL6 mice were mixed at various ratios (Figure 3b **bottom**), and further split into two aliquots for DNA extraction or antibody staining. Chimerism was analyzed from the extracted DNA by ddPCR and compared to chimerism measured by FCM (Figure 3b **top**). The measurements from both samples were nearly identical, with an R^2^ value of 0.998. This validates the assay for accurate detection of blood chimerism.

**Figure 3 legend.**
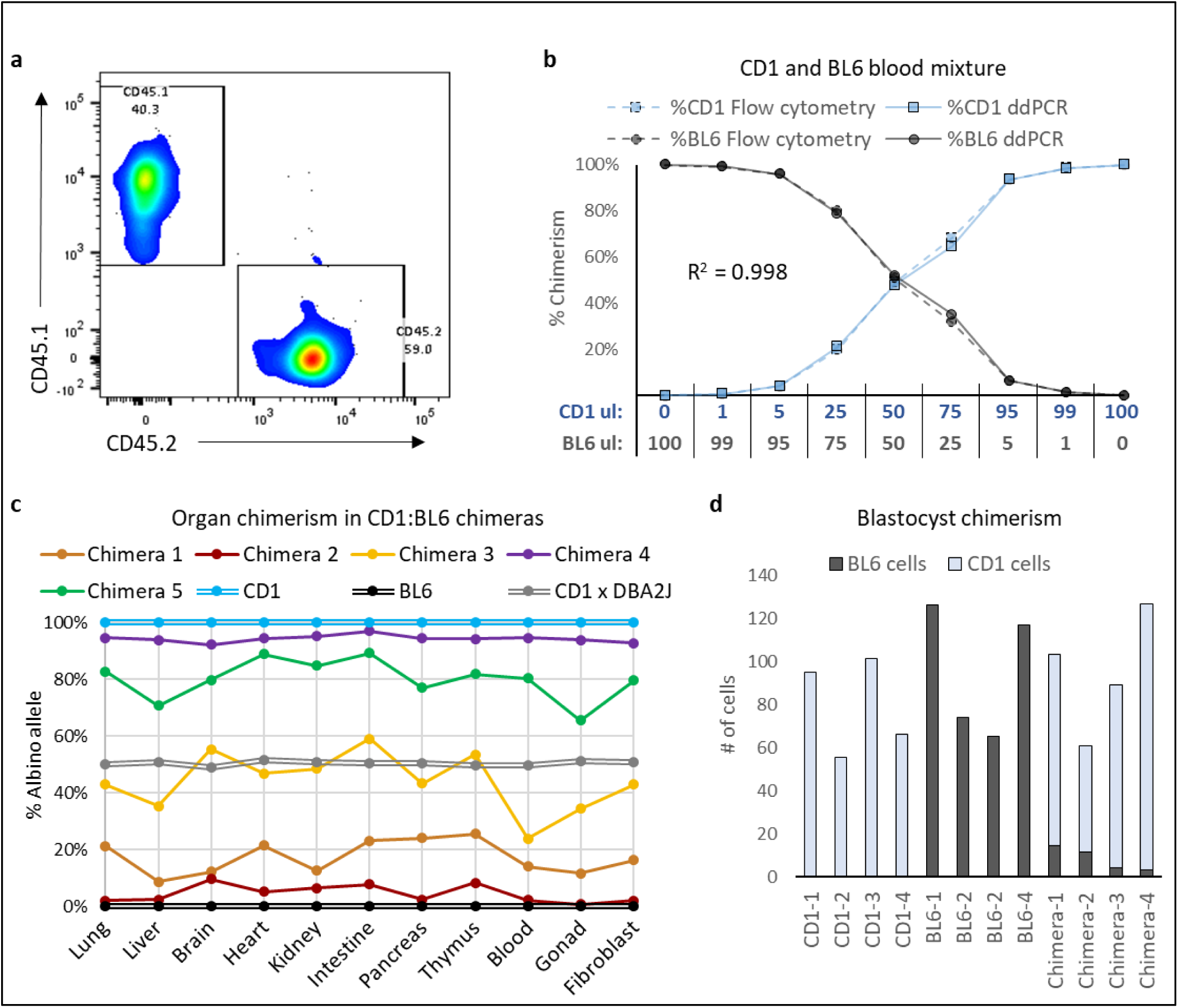
Validation in various organs and tissues. **a**. Flow cytometry plot of a CD1 and BL6 mouse peripheral blood mixture stained with CD45.1-PE-Cy7 (y-axis, CD1 strain) and CD45.2-BV421 (x-axis, BL6 strain) antibodies. **b**. Bottom: CD1 and BL6 blood mixed at different volumetric ratios. Top: Measured chimerism of CD1 and BL6 blood mixtures. Measured by FCM using CD45.1/CD45.2 antibodies (dashed lines) or ddPCR SND tyrosinase assay (solid lines). **c**. Chimerism measured with crude lysate of various organs from 5 chimeric mice (CD1:BL6 chimeras). CD1, BL6 and CD1xDBA2J F1 hybrid are non-chimeric controls. **d**. Number of BL6 and CD1 cells in blastocysts (1 lysed blastocyst per reaction). 4 CD1, 4 BL6 and 4 chimeric blastocysts (CD1 injected with BL6 embryonic stem cells) were analyzed.

### Organ/tissue chimerism: Analysis with direct lysis buffer

Purification of DNA from many samples can be tedious. Analysis of DNA directly from lysed cells without further purification would streamline the SND-ddPCR assay’s pipeline. To determine if crude cell-lysate from organs can be directly run on ddPCR and analyzed for chimerism, individual organs were harvested and digested with a tissue lysis buffer. The organs were harvested from CD1:BL6 chimeras (n=5) and controls (n=3), and the crude-lysate loaded directly into the ddPCR reaction. As expected, the degree of overall chimerism varied significantly from mouse to mouse, ranging from 6% to 96%, however chimerism in the organs within the same mouse was relatively stable (Figure 3c). The CD1 and BL6 controls were only positive for their respective alleles. A CD1 x DBA/2J F1 hybrid was also correctly 50% *Tyr*^*Alb*^ and 50% *Tyr*^*WT*^ in all organs, thus the extraction technique and use of crude lysate does not bias the results. These data suggest that SND-ddPCR assays can be used to accurately measure low to high chimerism in all organs, without the need for extracting clean nucleic acid.

One of the earliest and smallest structures during development is the blastocyst. To determine if the SND-ddPCR assay could detect chimerism in these small structures, individual blastocysts were lysed and analyzed as a single blastocyst per reaction. Four CD1 and four BL6 control embryos were positive for only the *Tyr*^*Alb*^ and *Tyr*^*WT*^ allele respectively (Figure 3d). In contrast, four CD1 embryos injected with a few BL6 mouse embryonic stem cells (mESCs) were all positive for both *Tyr*^*Alb*^ and *Tyr*^*WT*^ (all positive measurements were at or above the LoD). Additionally, since the entire blastocyst was loaded in one reaction, we could quantify the number of cells. There was an average of 90 cells/blastocyst, which is in agreement the expected number of cells in a late blastocyst. Therefore, the SND-ddPCR assay can be used to determine chimerism and overall number of cells in small structures that could not be readily analyzed by FCM, weight, and other methods.

## DISCUSSION

Robust quantification of chimerism is critical to measuring experimental outcomes. Here, we extensively validate a novel chimerism detection method that can be immediately implemented for many mouse strains and experimental workflows. We also showcase optimization and validation strategies that can be readily adapted to other assays. Since the method is a genetic assay, it has many attractive qualities, including (1) ability to distinguish a single nucleotide difference among otherwise identical cells, (2) transgenic modifications are not required, (3) agnostic to gene silencing, (4) resistant to tissue/cell-type dissociation bias, and (5) streamlined workflow.

One of the most powerful and unique capabilities of dPCR is the ability to robustly quantify allelic ratios that differ by only a single nucleotide. Most cells acquire point mutations during division, thus even subclones from the same parent line can be distinguished. Although we used a SND assay to identify a common SNP among different mouse strains, SND assays could be theoretically be applied to any SNP, including mosaic and cancerous genotypes(Pender et al., 2015). The ddPCR-based SND assay differs from other SND assays that rely on mismatches in the primer at the 3’-end. First, primer-mismatched assays require two separate reactions. Second, the polymerase will eventually extend the mismatched primer, even if the mismatch is in the 3’ end(Huang et al., 1992). Once this occurs, the PCR will continue with normal efficiency and a false-positive signal will be detected. Finally, the ultra-quantitative nature of ddPCR doesn’t just report the presence or absence of a SNP, but also reliably reports the concentration of each allele, nearly independent of overall concentration.

Use of a single, PCR-compatible lysis buffer can streamline detection of hundreds of samples. This can be performed in a high-throughput format (e.g., 96-well plate) with less hands-on time and minimal attention to individual samples. In contrast, cell dissociation for FCM requires tissue-specific and optimized lysis conditions, individual focus on each sample, and multiple spin/wash steps(Mitra et al., 2013). Additionally, most FCM samples need to be analyzed the same day before the cells die. Because of the easier and more reliable workflow, we have transitioned many of our chimerism analyses across organs from FCM to ddPCR(Nishimura et al., in press).

The dPCR detection strategy is the most robust and quantitative method compared to other PCR-based approaches. This is because the quantification strategy for dPCR does not require consistent or efficient amplification, but instead relies on simply counting the number of positive microreactions in individual clusters. dPCR is therefore more resistant to inhibitors and results in absolute quantification without references, as opposed to the relative quantification used in qPCR. However, even dPCR can be overwhelmed with inhibitors, preventing amplification. Although we could directly analyze crude lysates from most solid organs and tissues with the SND-ddPCR assay, it did not work well for whole blood. This was easily remedied with a hemolysis step, which is usually performed prior to FCM analysis as well. An alternative is to analyze a smaller volume of blood, however this results in loss of sensitivity.

When using a genetic assay to determine chimerism, ploidy must be considered. Liver, placenta, and cancer cells are often polyploid, which could lead to an overestimation of cell number(Celton-Morizur and Desdouets, 2010; Zybina and Zybina, 2020). Similarly, rapidly dividing cells spend a large proportion of time in S and G2 phases of the cell cycle, during which they have additional copies of chromosomes. In contrast, mature gametes are haploid, and red blood cells (RBCs) lack a nucleus, minimizing or precluding their detection respectively. However, except for RBCs, most changes in ploidy will only lead to a calculation error within a factor of 2, which is similar to the uncertainty associated with qPCR(Love et al., 2006). In addition, the lack of nucleus in RBCs is mostly advantageous, because it diminishes a blood measurement bias that would otherwise occur in heavily vascularized organs.

Perhaps the most frustrating property of ultra-sensitive assays is the potential for false positives. Since PCR can detect a single molecule, trace amounts of carry-over from other reactions can lead to false positives. Although the number of false positive droplets for each SND-ddPCR reaction is low and below our LoD cut-off, 29% had at least one false positive droplet (Figure 2d). In our lab, we occasionally PCR amplify and sequence the tyrosinase allele. Therefore, the false positive droplets in the SND assay could be due to post-PCR contamination from other experiments(Hu, 2016). In agreement, *Tyr*^*Alb*^ had a higher false positive rate than *Tyr*^*WT*^, likely because we more frequently amplify and sequence the albino allele. Therefore, it is recommended that each lab determine their own false positive rate and take precautions to prevent cross-experiment contamination.

It is important to understand the limits and confidence intervals of analytical platforms. We established our LoQ with >95% confidence that the measured value is within a factor of 2 of the true value. However, this is dependent on the volume of sample analyzed. Our confidence intervals were calculated assuming there are 17000 partitions (our average number of droplets per reaction), with a droplet volume of 0.85 nl (determined by BioRad). This corresponds to a total volume of 14.5 ul analyzed per reaction. Although at least 20 ul is usually prepared for each reaction, approximately 25% is lost during the partitioning process. Therefore, if the droplet volume or number of droplets decreases, the precision and sensitivity also decrease, and confidence intervals should be reassessed. This is most important near the lower limits of detection/quantitation and becomes negligible as concentration increases.

The SND-ddPCR assay accurately measured chimerism in all organs that were homogenized with lysis buffer and analyzed using the crude lysate. The hybrid control, which had an exact 1:1 ratio of *Tyr*^*Alb*^ and *Tyr*^*WT*^ in all cells, accurately reported this ratio in the tested organs. This shows that there is no allele bias that occurs during processing. The diverse range of average chimerism among the different chimeras highlights the variability of these types of microinjection experiments, emphasizing the need for robust quantification to measure subtle changes. In contrast, the chimerism among various organs in a single mouse was quite similar. This is expected, because mESC engraftment occurs at an early stage, and the mESCs we injected should not have a developmental predisposition to any tissues. Thus, in these experiments, chimerism is established stochastically at early timepoints, and relatively evenly distributes to all tissues. Similar experiments could be performed to discover tissue distribution biases of different cell lines.

We demonstrated the unique ability for the SND-ddPCR assay to count the overall number of cells in a sample by lysing and directly analyzing entire mouse blastocysts. With small samples, it is often impossible to dissociate into single cells without significant sample loss. Thus, this assay can be used as a primary or supplemental approach to measure organ/tissue size in small chimeric samples.

Reliable and robust measurements are key to scientific reproducibility. By simplifying data acquisition, researchers will generate higher resolution datasets with greater fidelity. dPCR chimerism assays are relatively easy to develop and extremely flexible. In addition, they have similar single-cell limitations compare to FCM-based assays, with numerous advantages. Therefore, we believe dPCR is a critical tool that can be used to streamline and strengthen chimerism/mosaicism measurements, with high accuracy and precision.

## ACKNOWLEDGEMENT

We thank Dr. K.C. Chan, M. Rivera, and N. Kowahara for laboratory and administrative support. This work was supported by grants from CIRM (LA1_C12-06917; DISC2P-11662), NIH (R01DK127144; R21OD030009), and the Ludwig Foundation. F.P.S. is supported by the National Science Foundation Graduate Research Fellowship and the Pat Tillman Fellowship. T.N. is supported by Japan Society for the Promotion of Science (JP18K14602 and JP18J00499). A.C.W. is supported by the NIH (K99HL150218), Leukemia and Lymphoma Society (3385-19), and the Evans P. Evans Foundation. J.B. is supported by the International Postdoc grant from the Swedish Research Council (2017-00344) and the Assar Gabrielsson Foundation, Sweden.

## AUTHOR CONTRIBUTION

F.P.S. conceived the research, performed experiments, and analyzed data; T.N. performed experiments and analyzed data; A.C.W. performed FCM and analyzed data; M.H. performed experiments; J.B. collected DNA and analyzed data; H.N. acquired funding and supervised experiments. F.P.S. wrote the manuscript and all authors edited and approved the final manuscript.

## MATERIALS AND METHODS

### Mice

All animal experiments were approved by the Administrative Panel on Laboratory Animal Care at Stanford University (APLAC #29042). All mouse strains were purchased from The Jackson Laboratory or Charles River, and housed with free access to food and water.

### Column purified DNA

Genomic DNA was purified using the DNeasy Blood and Tissue Kit (QIAGEN, Hilden, Germany) in accordance with the recommended protocol. For SND-ddPCR assay optimization (CD1 and BL6) and mouse strain genotyping (CD1, FVB, NSG, BL6, DBA2J, CD1xDBA2J), DNA was extracted from 2-4 mouse ear punches (~2 mm in diameter). Tissue lysis and homogenization was achieved after 3 hours at 55 °C followed by pipet trituration. For genotyping the C3H strain, DNA was extracted from 1 million cells. For blood, 15-50 ul was collected and lysed in accordance with Qiagen’s protocol. All samples were eluted in 100 to 200 ul of elution buffer.

### SND-ddPCR assay

Each ddPCR reaction was prepared and analyzed with the QX200 ddPCR system (BioRad, Hercules, CA) in accordance with BioRad’s standard recommendations for use with their ddPCR™ Supermix for Probes (No dUTP) unless otherwise stated. All reactions were mixed to 25 ul and contained up to 5 ul of purified genomic DNA or crude lysate, from which 20 ul were loaded into the droplet generator. Primers amplified a region of the mouse tyrosinase gene (forward, mTyr-F/1, 1.8 uM AATAGGACCTGCCAGTGCTC; reverse, mTyr-R/1, 1.8 uM, TCAAGACTCGCTTCTCTGTACA), which differs by a single nucleotide between albino and non-albino mice. Two hydrolysis probes with different fluorophores were used to detect either the albino allele (mTyr-alb-P/1, 0.25 uM, FAM-cttaGagtttccgcagttgaaaccc-Zen/IowaBlack) or the BL6 wild-type allele (mTyr-wt-P/1, 0.25 uM, HEX-cttaCagtttccgcagttgaaaccc-Zen/IowaBlack) in a single reaction with the above primers. Primers and probes were from IDT (Coralville, IA); probes contained the fluorophore at the 5’ end, the IowaBlack quencher at the 3’ end, and an additional internal ZEN quencher. Fifty PCR cycles were run with 30-second melting at 94 °C and 1-minute (min) combined annealing/extension at 60 °C. The average number of partitions after droplet generation was 17,000. Data analysis was performed with QuantaSoft Analysis Pro (BioRad, Hercules, CA).

### Peripheral blood FCM analysis

Peripheral blood from CD1 and BL6 mice were collected and mixed at various ratios. Following red blood cell lysis, samples were split for ddPCR analysis (as described above) and flow cytometric analysis as described previously(Wilkinson et al., 2020). For flow cytometric analysis, samples were stained with CD45.1-PE/Cy7 (clone A20; BioLegend, San Diego, CA) and CD45.2-BV421 (clone 104; Biolegend, San Diego, CA), washed and then run on a BD FACSAriaII, with data analysis performed using FlowJo software.

### Embryo culture and manipulation

Wild-type mouse embryos were prepared according to published protocol(Nagy et al., 2003). In brief, zygotes were obtained by oviduct perfusion from superovulated CD1 mice. Zygotes were cultured in KSOM-AA medium (CytoSpring, Mountain View, CA; K0101) for 2-3 days and blastocyst-stage embryos were collected for cell injection. For micromanipulation, mESCs were trypsinized and suspended in ESC culture medium. A piezo-driven micromanipulator (Prime Tech, Tsuchiura, Japan) was used to drill the zona pellucida and trophectoderm under microscopy and 5–8 ESCs were introduced into blastocyst cavities near the inner cell mass. After blastocyst injection, embryos were cultured for 1–2 hours. Mouse blastocysts were then transferred into uteri of pseudopregnant recipient CD1 female mice (2.5 days post coitum) to generate chimeric animals.

### Crude extraction of DNA from organs

To lyse and homogenize solid organs, mice were euthanized and a ~1.5 mm^3^ cube of the respective organ was added to 200 ul of lysis buffer. The lysis buffer contained 0.1% SDS, 5 mM EDTA, 1 mg/ml proteinase K (Thermo Fisher Scientific, Waltham, MA), 15 mM tris (pH 8), and 100 mM NaCl. Lysis was performed for 6-12 hours at 55 °C. The lysate was then homogenized with a pipet or 20 gauge hypodermic needle if needed, and all samples were heated to 80 °C for 10 minutes. For blood, 15-50 ul was collect and red blood cell lysis performed prior to addition of lysis buffer. Lysis was then performed for only 20 minutes at 55 °C, followed by 10 minutes at 80 °C. For all samples, debris was pelleted from the lysate by centrifugation at 6000 g’s x 3 minutes, and 1-5 ul of the supernatant analyzed by SND-ddPCR.

### Blastocyst ddPCR

CD1 and BL6 morulae were harvested as described above. Four CD1 morulae were injected with eight BL6 mESCs. The embryos were cultured until E4.5, after which they were transferred into 4 ul of lysis buffer in PCR tubes. The lysis buffer was similar to the buffer used for organs, except it contained only 0.2 mg/ml proteinase K. 6 ul of water was then added to each PCR tube, and the tubes were heated to 55 °C for 10 minutes and 80 °C for 10 minutes. The ddPCR reaction mixture was prepared directly in the tubes contained lysed blastocysts.

## SUPPLMENTAL MATERIAL

**Supplementary Figure 1 legend.**
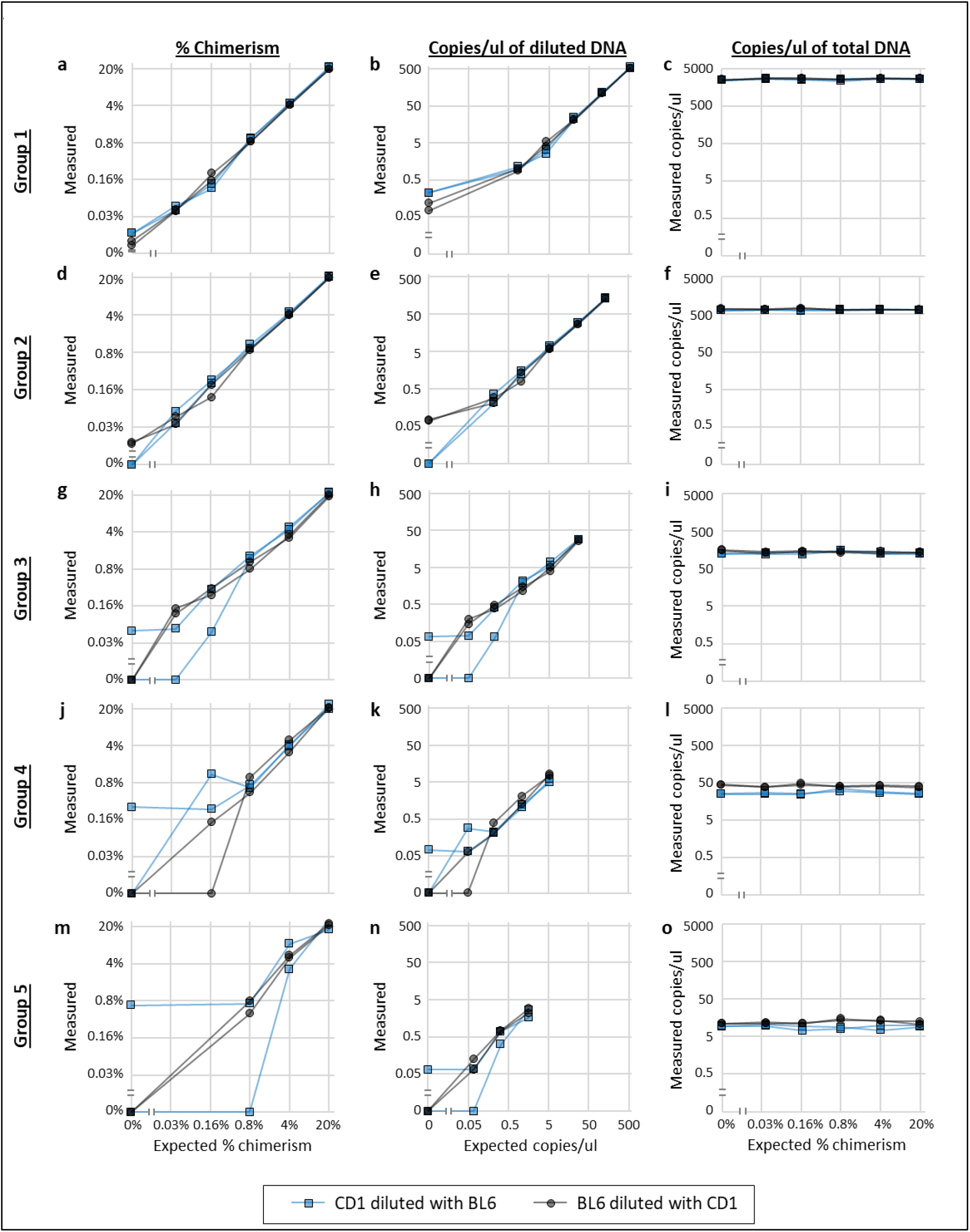
CD1 and BL6 dilution matrix. Measured abundance of Tyr^Alb^ and Tyr^WT^ allele in mixtures of CD1 and BL6 genomic DNA; measured results compared to expected results from dilution matrix. Axes are log-scaled. Blue boxes show two separate measurements when Tyr^Alb^ is the low abundant allele (CD1 diluted with BL6). Blue lines connect a prepared dilution series within the group. Gray circles show two separate measurements when Tyr^WT^ is the low abundant allele (BL6 diluted with CD1). Gray lines connect a prepared dilution series within the group. The left column (**a, d, g, j, m**) plots the measured %chimerism of the low abundance allele in Group 1-5 respectively compared to the expected %chimerism. The middle column (**b, e, h, k, n**) plots the measured concentration of the low abundance allele in Group 1-5 respectively compared to the expected concentration. The right column (**c, f, i, l, o**) plots the measured total concentration (sum of both alleles) in Group 1-5 respectively for each sample within that group; they are the same in each group as expected.

